# Network analysis of marmoset cortical connections reveals pFC and sensory clusters

**DOI:** 10.1101/2024.03.04.582858

**Authors:** Bernard A Pailthorpe

**Affiliations:** Brain Dynamics Group, School of Physics, University of Sydney, Sydney, NSW, Australia

**Keywords:** marmoset, structural connectivity, network, module, cluster, hub, pFC

## Abstract

A new analysis is presented of the retrograde tracer measurements of connections between anatomical areas of the marmoset cortex. The original normalisation of raw data yields the fractional link weight measure, FLNe. That is re-examined to consider other possible measures that reveal the underlying in link weights. Predictions arising from both are used to examine network modules and hubs. With inclusion of the in weights the Infomap algorithm identifies eight structural modules in marmoset cortex. In and out hubs and major connector nodes are identified using module assignment and participation coefficients. Time evolving network tracing around the major hubs reveals medium sized clusters in pFC, temporal, auditory and visual areas; the most tightly coupled and significant of which is in the pFC. A complementary viewpoint is provided by examining the highest traffic links in the cortical network, and reveals parallel sensory flows to pFC and via association areas to frontal areas.

## Introduction

Our early knowledge of brains came from tracing neural pathways. Modern network science offers a computational equivalent that can reveal local structures worthy of further study. The marmoset brain has well developed frontal lobes, so provides an accessible system for studying cortical function. In evolutionary terms it is midway between mouse and higher primates, so provides a good entry point for subsequent modelling. The Rosa lab has accumulated significant data from tracer injections and labelled cell counts to provide a measure of link weights in the marmoset cortex [Majka et. al. 2016, 2020]. That study sampled 55 injection sites chosen amongst 116 anatomical areas. They also conducted a detailed network analysis of the fully connected subnetwork of 55 nodes [Theodoni et. al. 2022]. The present study attempts an analysis of the partially sampled network of all 116 areas of the cortex. While incomplete it does include valuable additional data. Given that the choice of the 55 injection sites was guided by expert biological insights, the available data is likely to capture a significant fraction of the important links in the cortex.

There is also an an opportunity to apply methods used previously for worm, mouse retina, mouse, rat and macaque to study this system. Various groups have used different measures of connectivity, or link weights, depending on the experimental techniques used (EM, tracer, MRI) and the research questions examined. Thus synaptic contact areas (mouse retina), tracer volume and raw cell counts (mouse brain) and fractional measures (macaque and marmoset cortex) have been used. Consistency with prior research is one reason for a given choice, while the merits of the varying approaches are still being assessed. The analysis methods herein follow those used for the mouse retina [Pailthorpe 2016] and the mouse brain [Rubinov, et. al. 2015, Pailthorpe 2019], and are compared with other choices.

The marmoset network data is at mesoscale, describing links between anatomical areas. Variability of repeated tracer injection volumes suggested Fraction of Labelled Neurons, extrinsic, FLNe as a practical measure of link strength, as used on retrograde tracer data from macaque cortex [Markov et. al. 2011, 2014], and as originally reported for the marmoset data used herein. The present study uses the more extensive, recent marmoset data [Majka et. al. 2020] to renormalise that fraction to a more direct measure of source-target link weights based on LNe, the number of Labelled Neurons, extrinsic. This is consistent with the mouse brain tracer data [Oh et. al. 2014] that reported links weights as raw connection strength, CS [Rubinov, et. al. 2015], specifically the measured volume of fluorescent tracer detected in a linked target area. That should be proportional to the number of labelled neurons – essentially equivalent to LNe in the present notation – a question examined herein. The benefit of this approach is that it discriminates varying in link weights to target nodes, and then incorporates those weights in the subsequent analysis.

It is illustrative to begin the network analysis with counts of in and out links to/from nodes, i.e. the node degree, followed by summing weights of in- and out-links to find the weighted in- and out-degree of each node (node strength). While elementary, revisiting those measures reveals consistencies within the dataset and with other studies, and suggests the new approach adopted herein. The distributions are variously exponential, normal or log normal, depending on the species, methods used, and scale of the measured areas. The present analysis of the marmoset data also covers: dependencies of the basic link weight measurements on injection and target volumes that facilitates a re-analysis of the raw data using a rescaled measure of links weights (cf. Methods, Supplementary Material); link weight distribution; link weight-distance plots to confirm exponential decay; modular decomposition using InfoMap and Louvain methods, and identification of network hubs; and tracing evolving links around hub nodes to identify local clusters and pathways in the cortex, along with sensory pathways. Link tracing around key hubs reveals local clusters in pFC, auditory, association and visual areas; the pFC cluster is the most tightly interlinked. Visualisation of pathways around hubs and connectors, and those hosting high network traffic, suggest parallel pathways from auditory and visual sensory areas to pFC and motor areas. By contrast modular decomposition based on the fractional measure FLNe, using both InfoMap, herein (cf. Supplementary Material), and Louvain methods [Theodoni et. al. 2022], gives prominence to the visual areas.

## Methods

Mesoscale connectivity data for the marmoset cortex is available [Majka et. al. 2016, http://marmosetbrain.org] from retrograde viral tracer injections to 55 target areas selected from amongst 116 anatomical areas of the cortex. This data is for ipsi-lateral links in the left hemisphere. Characteristics of the original experiments have been reported in detail [Majka et. al. 2020, Theodoni et. al. 2022], and the basic data is summarised in the Supplementary Materials. The marmoset link weights were reported as Fraction of Labelled Neurons, extrinsic, FLNe – a measure also used for earlier macaque data [Markov 2011, 2014]. The use of FLNe means that that the total weight of all in-links to a given node is 1, or 0. Even so the original data set still shows the variation in the number of in links to each node, displayed in Figure S1. The weighted degree, or node strength, distributions are shown in Figure S2. The out-degree appears to have an exponential decay, as observed for other species, eg. the nematode worm C. Elegans [Varshnay et. al. 2011], mouse retina [Pailthorpe 2014] and mouse brain [Oh et. al. 2014, Pailthorpe 2019]. However the original weighted in-degree adopts only values of 0 or 1 since fractional measures sum to 1 for each node, which follows from the definition of FLNe. That factors out differences of in link weights between nodes, so that some network information is discarded.

### Rescaled link weight data, LNe

In this analysis the original FLNe data is rescaled to be consistent with the mouse data [Oh et. al. 2014] and to uncover the underlying in link weights. This is possible with the more recent, larger set of measurements (Majka et al, 2020] that includes sufficient detail to enable the fractional normalisation, used above, to be recalculated. Here a cortex wide, common normalisation for all nodes (representing anatomical areas) is adopted so that the underlying weights of in links can be recovered. The procedure is detailed in the Supplementary Materials. The new data uses the same 55 target nodes as the original data, multiple tracers, with repeated experiments, so 143 experimental results are available. Extensive raw experimental data was reported: ie. each injection volume, number of Labelled Neurons (LN) both intrinsic (to the injected volume) and extrinsic (meaning all source neurons), and their total: LNi, LNe, LNtot respectively. This extra data enables the normalisation, that was used to calculate FLNe, to be recovered, and to produce the underlying link weights. That normalisation factor varies for each target node since it is a measure of the weighted in degree, or node strength. Dependencies of LNi and LNe on injection volumes and node volumes are explored in Figs. S5 and S6 and analysed in the Supplementary Material, and informs the choice of link weight measure.

Here the direct measure, LNe, is used so that the underlying weight distribution for the in-links is revealed. LNe is the number of labelled neurons, extrinsic (ie. not in the target area, and including all those detected in related source areas). This is consistent with the mouse brain tracer data [Oh et. al. 2014] that reported links weights as raw connection strength (CS), specifically the measured volume of fluorescent tracer in a linked target area. That should be proportional to the number of labelled neurons – essentially equivalent to LNe in the present notation. A caveat for that comparison is that it assumes uniform and constant neuronal density: that has been measured for Marmoset [Atapour and Rosa 2019] and exhibits ∼10% variation between some areas, and an Anterior-Posterior gradient.

In network terms the measured LNe is equal to the weighted In Degree (or node strength) of the injected target node: this can be calculated both for out links (sum over linked targets) and for in links (sum over linked sources). In an explicit notation, the reported FLNe(s, t) is the fractional link weight from source s to target t. Similarly LNe(t) identifies the injection site for the retrograde tracer with the target node. This LNe(t) is the total weight of incoming links from linked source nodes: i.e. LNe(t) = Σ_s_ LNe(s, t), where the sum is over all linked source nodes (s). That sum is just the weighted In Degree of the target node (cf. Fig. 1). It varies across nodes (i.e. areas) while, by contrast, the corresponding fractional measure FLNe(t) is either 0 (no incoming links), or 1 (target linked from one or more sources); cf. Fig. S2. The array of LNe(s, t) is just the network adjacency matrix, attached at Supplementary Data, which forms the basis of subsequent calculations. The detailed procedure for its calculation is presented in Supplementary Materials.

**Figure 1.**
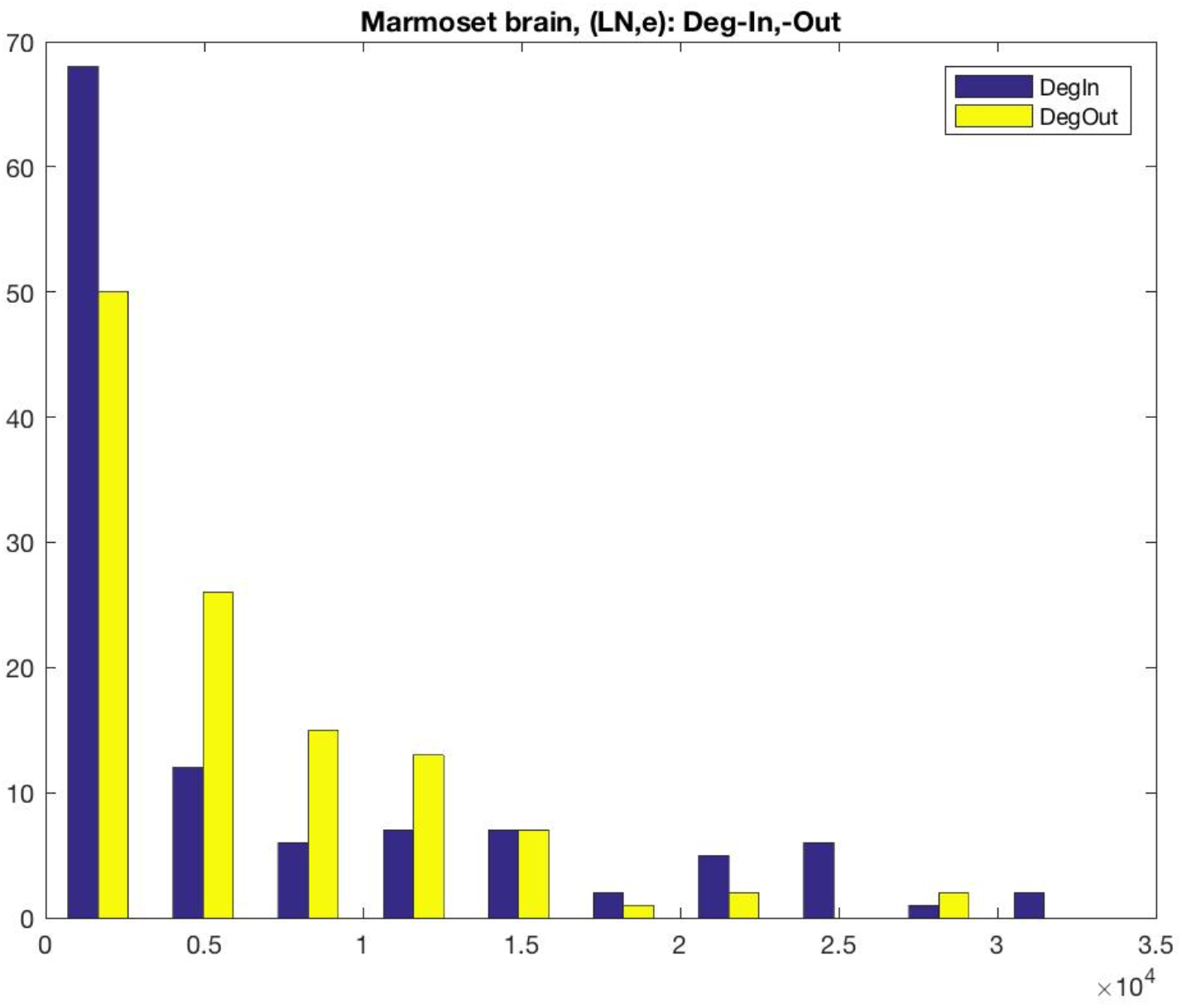
Distribution of the weighted in- and out-degree z(or node strength), calculated using the rescaled measure LNe, for anatomical areas of the marmoset brain.

### Network modules and hubs

Here two methods are used to decompose a network into modules, that have more linkages between nodes within a module than between them. Results are reported using both data sets, based on FLNe and on LNe, as discussed above. First, the InfoMap method and software models random walks over the network [Rosvall and Bergstrom 2008] and optimises an information theory based metric [Rosvall and Bergstrom 2007]. Second, the Louvain method [Blondel et. al. 2008] agglomerates nodes and then re-iterates to optimises the modularity metric [Newman 2003]. They generally produce similar results despite being conceptually quite different algorithms.

A related analysis, originally applied to metabolic networks within a cell [Guimera and Nunes Amaral 2005], leads to identification of network hubs. Two measures are calculated and plotted together: the participation coefficient of node i, P_i_, is a measure of the fraction of a node’s links within a module, compared to all its links. The limit P_i_ →1 indicates that the node has wide ranging links between modules, while P_i_ →0 indicates that the nodes links are primarily local, within its own modules. The second measure, the z-score, z_i_ indicates how well connected a node is to other nodes in its own module (a membership score, or within-module degree measured in standard deviations from the mean). These measures can be calculated separately for in-links and out-links. 2D plots of z_i_ vs. P_i_ have been divided into regions that classify nodes and types of hubs, as illustrated in the figures below. The regions in the 2D plot and the hubs’ classifications follow the heuristics developed for metabolic networks [Guimera and Nunes Amaral. 2005] to describe classes or roles assigned to network nodes. The important ones are: Role 6 (R6), connector hubs (many links between modules); R5, provincial hubs (links preferentially within module); R4, non hub kinless (links across most, or all, modules); and R3, non-hub connectors (many links to other modules). A related analysis has been applied previously to cat and macaque brain networks [Sporns et. al. 2007]. That study used a binary connection matrix, i.e. unweighted links. For comparison the module decomposition computed using marmoset FLNe is presented in Table S1 and discussed in the Supplementary Material. The difference in in-weights in the two methods produced different module decompositions.

The key measure that separates hubs from non hubs is the z score. That measures the node’s within-module degree, relative to the average and normalised by the standard deviation (of within-module degree, of all nodes in that module). The cut off originally assigned, in a cellular biology application, for classification as module hubs was z > 2.5 standard deviations above the average. In the present study (cf. Fig. 2) some nodes are more than two standard deviations above the mean, so credibly could be classified as marginal hubs – thus they could be investigated further.

**Figure 2.**
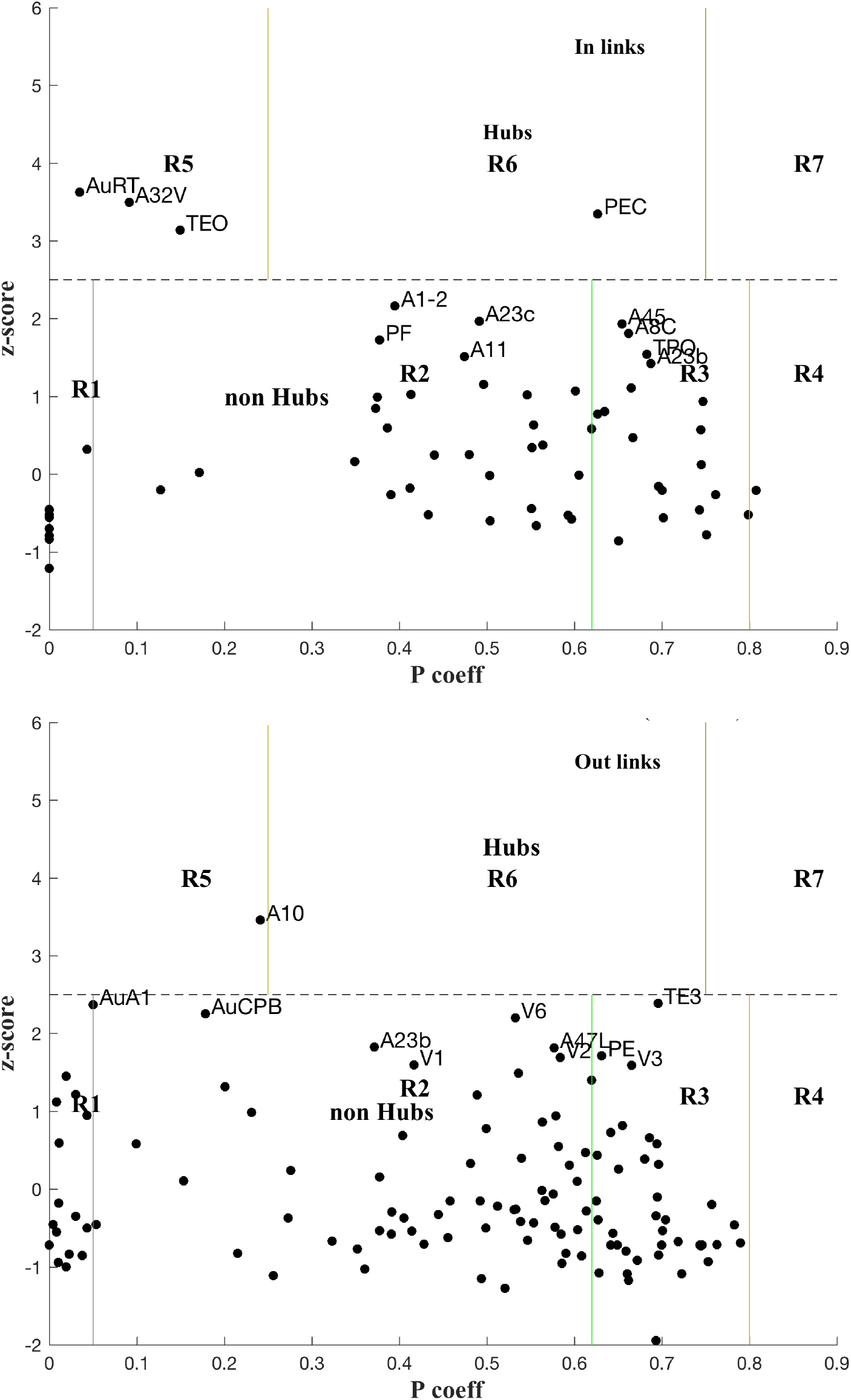
Plot of z-scores vs. participation coefficients for the marmoset cortex, calculated using on the LNe data (cf. Methods, Supplementary Material) for in-links (A. top panel) and out-links B. (bottom panel).

### Link Pathways

Having identified modules and key hubs and connector nodes analysis of their local link network can shed light of cortical organisation. To explore this the ideas implicit in temporal networks [Holme and Samarki 2012] were used to trace time evolving links to or from selected nodes. Assuming constant signal velocity, here taken to be 1 m/s, as an example typical of local circuits that may have unmyelinated axons, provides a scaling between link distances and time. Thus following links of longer pathlength also follows the time evolution of pathways. Link tracing covered all linked first nearest nearest neighbours and extended to second linked neighbours, allowing for a 2 ms synaptic delay (ie. equivalent to 2 mm extra pathlength). A Matlab code assembled all the links with a selected node[s], then resorted them into increasing link distances, or shells of neighbours, and incrementally drew the evolving network of local links in 3D. This facilitated visualisation of link patterns around the key hubs. Thus locally clustered nodes and their local and global links readily became apparent.

All calculations were performed with Matlab codes available from standard repositories (BCT at https://sites.google.com/site/bctnet/, www.mapequation.org, NIFTI tools: http://mathworks.com/matlabcentral/fileexchange/8797), or written Matlab codes lodged at https://github.com/BrainDynamicsUSYD/MarmosetCortex/.

## Results

Here a direct link weight measure, LNe, is employed so that the underlying weight distribution for the in-links is revealed. LNe is the number of labelled neurons, extrinsic (ie. not in the target area, and including all those detected in related source areas). The rescaled link weights, LNe are now in the range 0.03 – 10^4^. The resulting link weight distribution (log_10_ scale, Fig. S4) can be compared with the distribution using the original fractional measure FLNe (Fig. S3). The weighted in- and out-degree distributions are shown in Figure 1.

Both the weighted In- and Out-Degree distributions decay rapidly, in contrast to the results using FLNe (cf. Fig. S2). Plots of a fitted exponential distribution (Matlab fitdist tool; not shown) indicate that the in Degree falls off more rapidly than exponential, as also evident in Fig. 1, compared to the out Degree. Possibly some low weight in links are missing, since the in linkage data is incomplete with only 55 of the 116 anatomical areas being tracer injection sites. Here an extra 1611 links are included that these are out links from the non sampled areas to any of the 55 injection sites. The present results are more in line with tracer results for Mouse [Oh et. al. 2014, Pailthorpe 2019] and EM based results for worm [Varshnay et. al. 2011] and mouse retina [Helmstaedter et. al. 2014, Pailthorpe 2016].

The log link weight - distance plot (Fig. S7) indicates an exponential decay with large scatter, as observed for other species, and a linear fit yields weight ∼ e^-dist/4.57^ (R^2^ = 0.16, with distance in mm). For comparison a log-log plot similarly displays large scatter, and produces a scaling law: weight ∼ dist^-2.00^ (R^2^ = 0.23), compared to weight ∼ dist^-2.05^ found for mouse [Rubinov et. al. 2015]. The difference between the two fits is too marginal to discriminate. The inverse square law decay is consistent with geometrical embedding. For comparison, functional connectivity data in human brain produces a weight - distance decay midway between exponential and power law [Roberts et. al. 2016].

### Network modules and hubs, using LNe

The module analysis computed using marmoset FLNe is presented in Table S1 and discussed in the Supplementary Material. Module membership computed using the Infomap method with rescaled weights derived from LNe are summarised Table 1, with a full list attached at Supplementary Data. Probability flow is a key variable in the Infomap method and priorities nodes, and modules, based on the fraction of random walkers transiting a node – essentially its a measure of local network traffic.

**Table 1.**
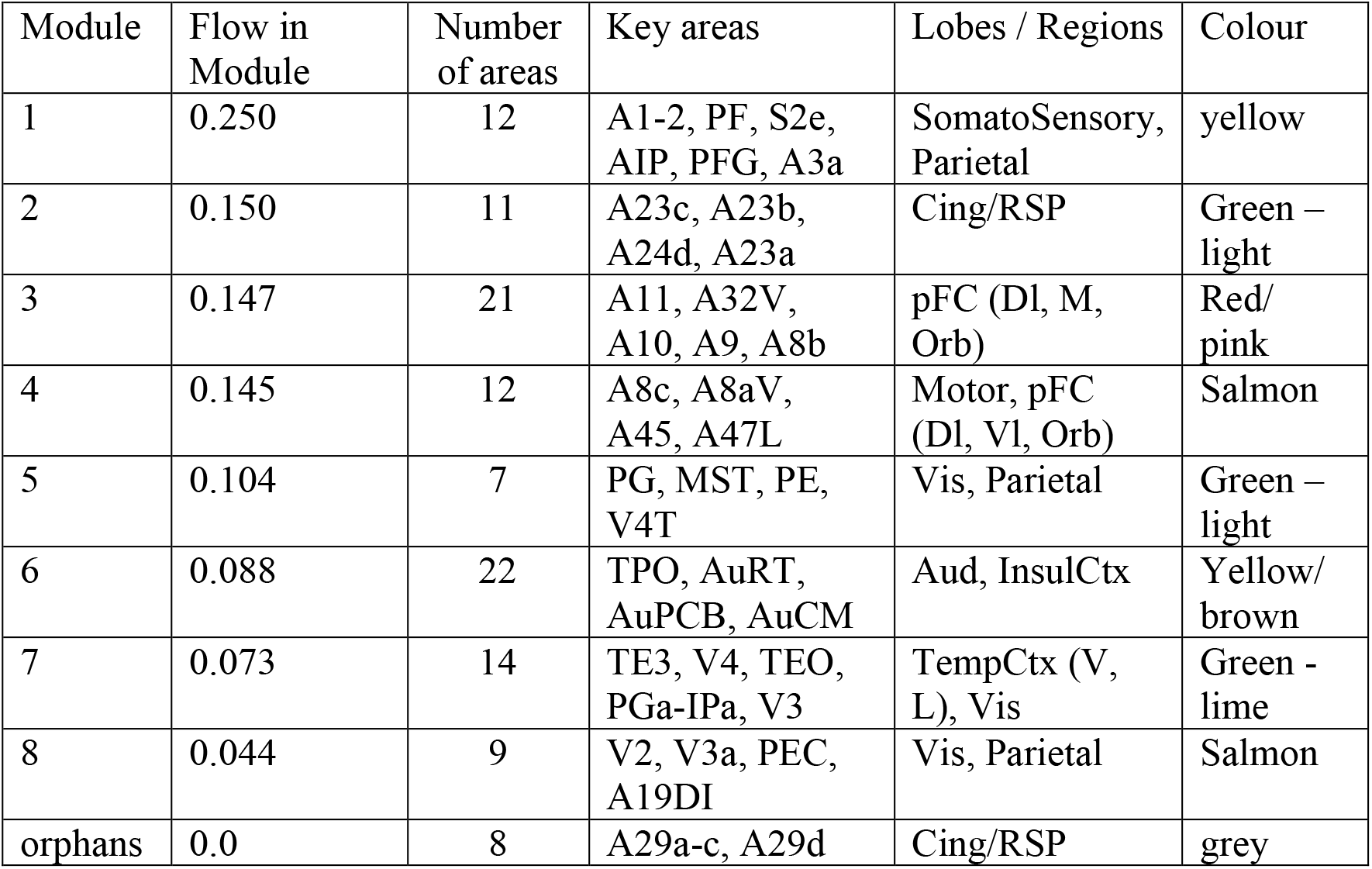
Infomap modules for marmoset cortex ipsi-lateral links, using the rescaled LNe measure of link weights. Key members are listed in order of probability flow; colour coding follows the Marmoset Atlas and is used in the figures.

Here the module membership is somewhat similar to that found with the original FLNe measure (cf. Table S2), however the order is changed, both within and between modules. Now the somatosensory system hosts the most network traffic, and here is separated from the motor system. The modules containing the largest number of anatomical areas are #3, associated with pFC and #6, auditory cortex. Note that this module assignment likely will be revised when in links are measured to the remaining 61 (ie. 116-55) targets are measured and reported. Here also the auditory system constitutes its own module, and the previous modules 6 and 7 (cf. Table S2) are merged into a single module (6). The visual system is present across three modules. The contrast between Tables 1 and S2 is that differences in the weight of in links are accounted for rather than being factored out as in the common normalisation forced by the fractional measure, FLNe.

For comparison the Louvain method [Blondel et. al. 2008], an agglomerative method, was also used to find the modular decomposition of the marmoset cortex, generating 7 modules that are closely aligned with the InfoMap results reported here. The modularity metric is Q = 0.52 [Newman 2003]. It reported no singletons (isolated nodes). Interestingly the Louvain method when applied to an unweighted adjacency matrix did not decompose the network into modules, indicating that the link weights play acritical role in resolving the component modules. A separate analysis of the full connected 55x55 connectivity matrix using the Louvain method [Liu et. al. 2020], with tuned sensitivity, also produced 8 modules and assigned prominence to frontal and auditory areas, in agreement with the present analysis.

Classification of nodes into types of hubs and connectors is illustrated in Figure 2 with plots of z_i_ vs. P_i_, the module membership score (within-module degree) and participation coefficient, respectively, of each node (cf. Methods), for both in and out links. The zones for each class of role are labelled. Hubs and nearby, possible candidates are also labelled. The boundaries were based on heuristics developed for cellular metabolic networks [Guimera and Nunes Amaral 2005], eg. z > 2.5 for hubs, and that may need to be revisited for cortical networks.

The rescaled link weights preserve differences of in links between various nodes, and now reveals in-hubs: PEC (Parietal) as a Connector in hub (R6), and AuRT (Aud), A32V (M-pFC) and TEO (Temporal) as Provincial in hubs (R5). These in hubs are missed in the analysis using FLNe (cf. Fig. S8). Three nodes, A1-2, A23c and A45, are close to the borderline for in-hubs, so might be marginally classified as hubs since the z-score cutoff (2.5) is somewhat arbitrary in that it was originally determined by a heuristic rule for another biological application [Guimera and Nunes Amaral 2005]. There are no out-link Connector Hubs (R6; cf. Fig. 2), but PGa-IPa might be considered as marginal; A10 is a Provincial out hub (R5), with AuA1 and AuCPB as marginally so. TE3 is a borderline connector out hub (R6). The full list of p, z values for the 116 nodes are attached at Supplementary Data.

For in links, four nodes are prominent non-hub connector nodes (R3; ie. well linked to other modules): A45, A8c, TPO and A23b; another 13 are also R3 but less well linked: PG, V4, TE3, A47L, PGa-IPa, A23a, V4T, AIP, PFG, A8aD, A6M, LIP and V5. Note that there are commonalities with the lists using FLNe (Supplementary Material), but also differences. For out-links six nodes are prominent non-hub connector nodes (R3): PE, V3, A8aV, PG, V4 and TPO - again in order of decreasing P; another 22 nodes are classified as R3, but are less well connected. One node for which in links were measured (ie. an injection site) is classified as peripheral (R1) for in-links: A32V, along with eight nodes that were not measured for in-links: AuAL, AuR, A25, AuRTL, AuRM, TPro, AuRTM and APir. As a check these network based classifications need to make sense biologically.

The top 8 hub nodes are: A10, TE3, AuA1, AuCPB, AuRT, A32V, TEO and PEC. They are plotted in 3D with their in and out links in Figures 3 and 4, respectively. Only links of weight >1 and length < 5mm are shown for clarity, and to reveal the local clustering. The very weak 135 out- and 51 in-links, of weight < 1, are omitted to avoid clutter. Colours follow the marmoset Atlas [Paxinos et. al. 2012]; a grey underlay is introduced to provide some contrast, which may shift the colours.

**Figure 3.**
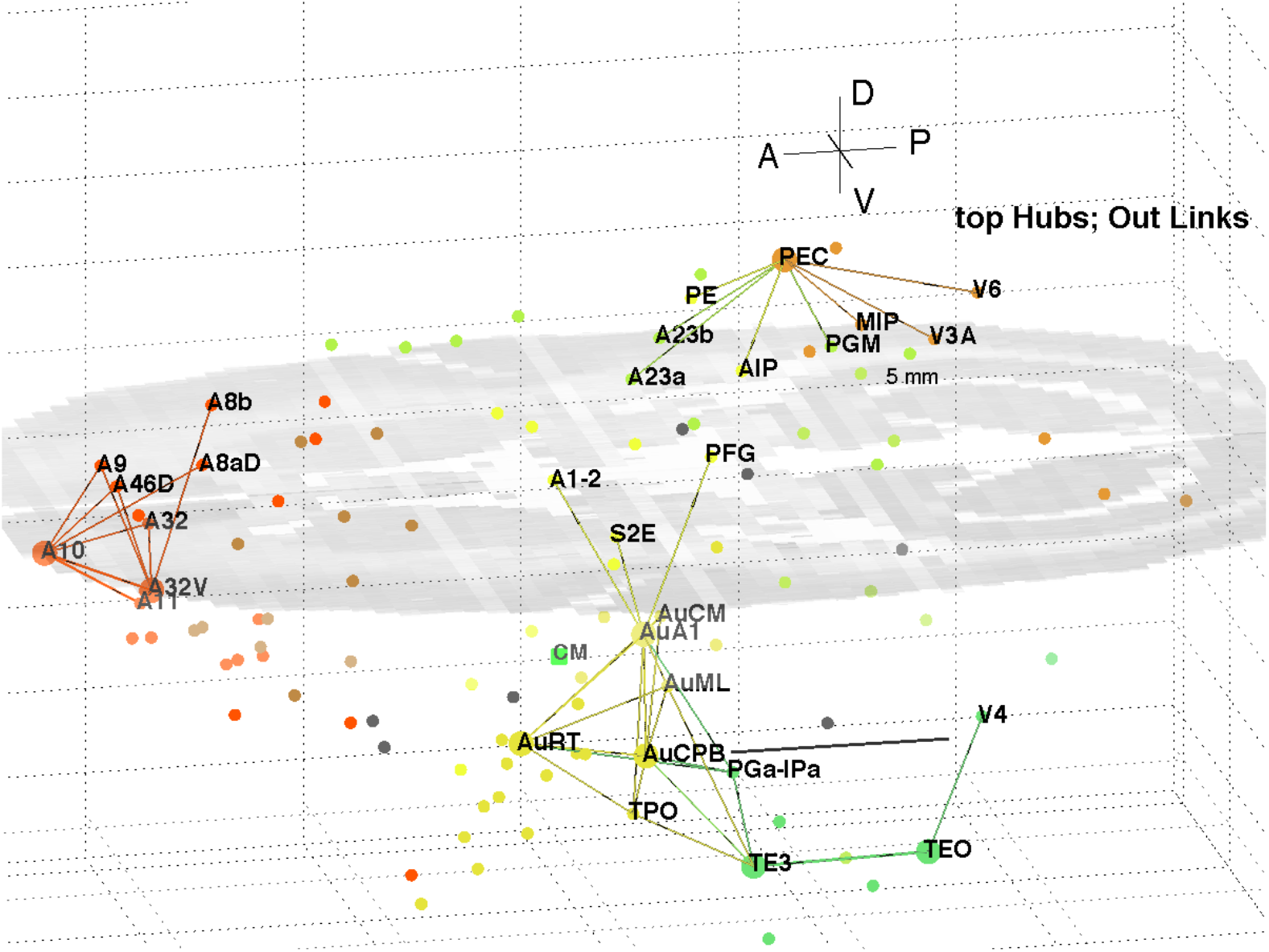
3D plot of network hubs and their in links of length < 5mm. Nodes are coloured by their module membership (cf. Table 1) and links by the source node membership. The mid level horizontal plane image is a slice from the marmoset cortex volume image to aid perspective, along with the background grid. Ipsi-lateral links for the left hemisphere were reported. The background grid is 5 × 2 mm and the A-P scale bar (near V4) is 5mm.

**Figure 4.**
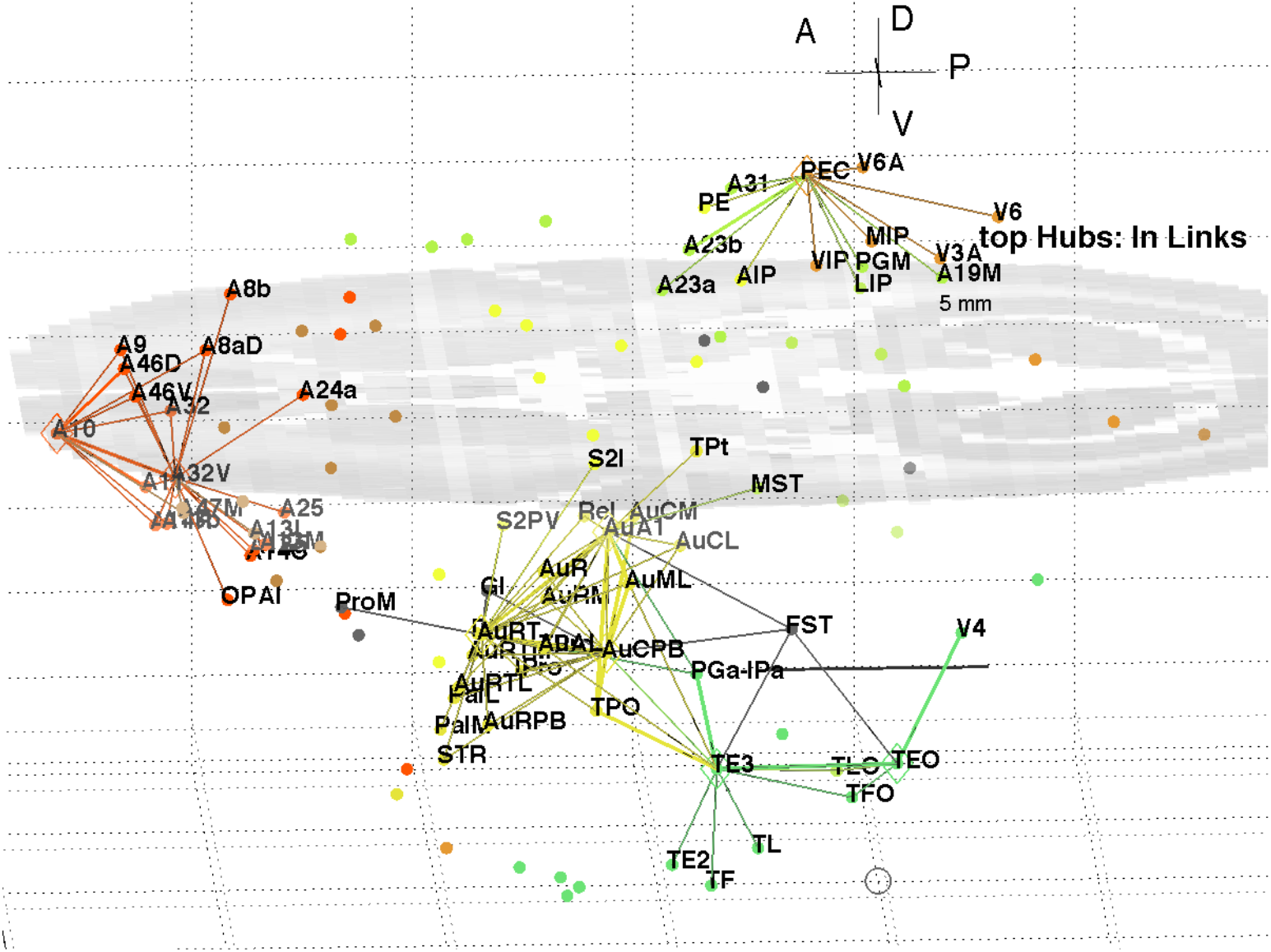
3D plot of network hubs and their out links of length < 5mm. Nodes are coloured by their module membership (cf. Table 1) and links by the target node membership. Other details as in Fig. 3.

A separate analysis of the 55x55 subnetwork using participation coefficients only [Liu et. al. 2020] identifies frontal, auditory and association areas as most hub like, followed by visual areas, and singles out A10 as having unusually large out strength, in agreement with the present study.

Continuing the path tracing of Figs. 3 and 4 to longer distances (∼ 10 mm) reveals direct links from the auditory areas AuCPB and AuA1 to the frontal areas; and bridging nodes between the visual and auditory sensory areas to be PEC, TEO and TE3; with PEC later on linking to motor areas

Similarly, the top 9 connector nodes are: PE, V3, A47L, V6, A4b, A45, A1-2, and A23c. They are plotted in 3D, along with their close in and out links in Figures 5 and 6, respectively. These show the evolving fabric of links between the hub clusters.

**Figure 5.**
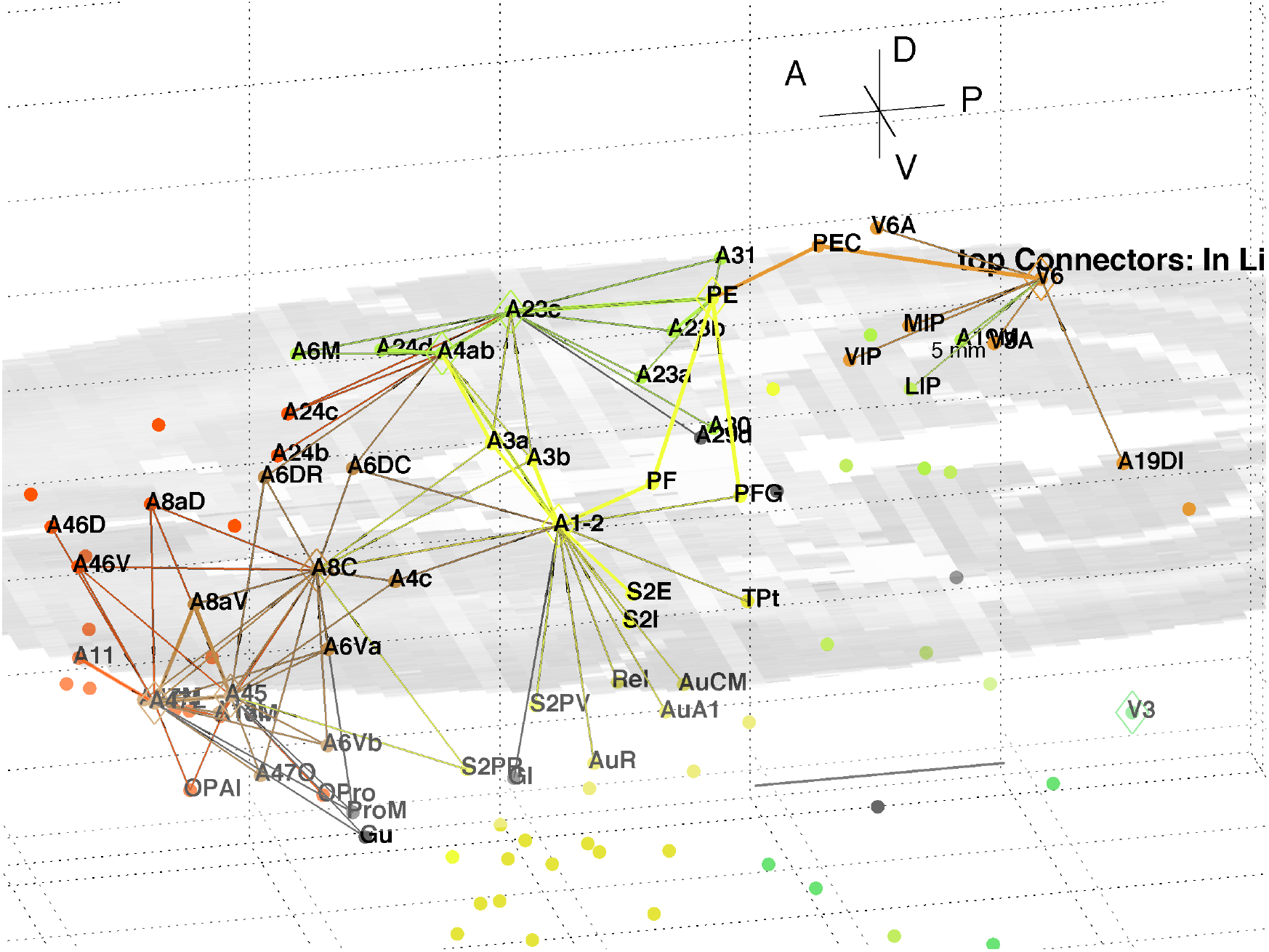
3D plot of network connector nodes and their in links of length < 5mm. Nodes are coloured by their module membership (cf. Table 1) and links by the source node membership. Other details as in Fig. 3.

**Figure 6.**
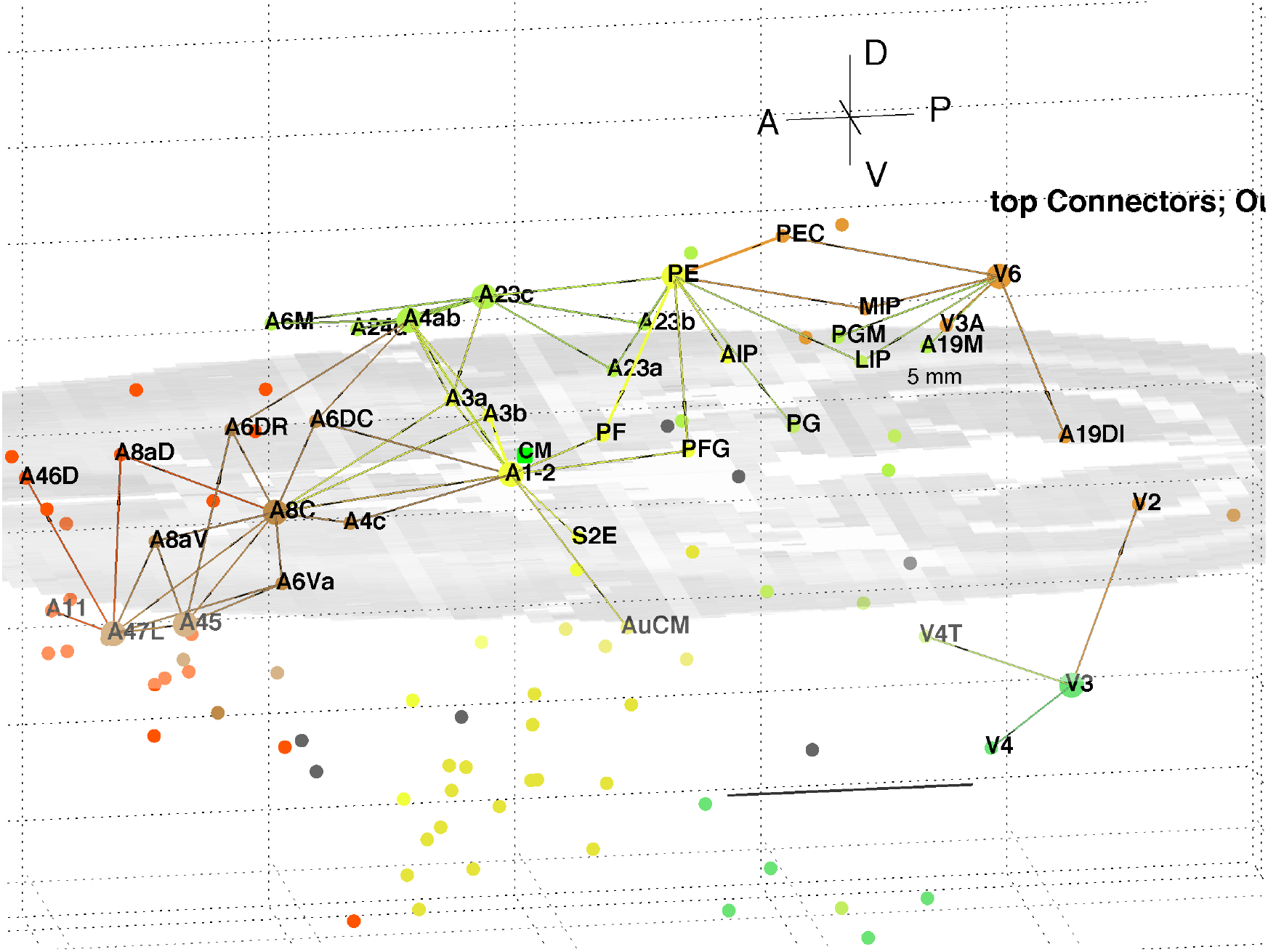
3D plot of network connector nodes and their out links of length < 5mm. Nodes are coloured by their module membership (cf. Table 1) and links by the target node membership. Other details as in Fig. 3.

Together these visualisations show the major local clusters and their interlinkages in marmoset cortex. Sensory Pathways were traced (cf. Methods) from visual and auditory cortex to anterior cortical areas. Aside from relatively weak direct links (eg. V2 – A8aV, V2 – A32) high weight links to the frontal areas from visual areas requires two steps, via intermediate transit nodes: eg. V2 via V4, V5, MST, TEO, TPO, and Opt. Those variously on linked to areas of the pFC cluster: A32, A46D, A10, A11, A32V and A9. For the auditory areas direct links are from AuA1 via TPO to the hubs A32V and A11, and from AuCPB direct to A10, A32V, A11 and A46D, all targets in the pFC cluster.

### High traffic links derived from LNe

The InfoMap analysis also yields the probability flow through individual nodes which, in turn, allows calculation of flow over links. This flow is the fraction of random walkers that pass over the node or link. That allows identification of the highest signal traffic links in the whole cortex, as displayed in Figure 7. It displays the top 200 of both in and out links, being the top 15% highest probability flow amongst all cortical links. Those linkages follow a similar pattern to that derived via hubs and connectors as shown in Figs. 3-6. The colours identify module membership (cf. Table 1) and follow those of the marmoset Atlas. The mid level horizontal plane image is a slice from the marmoset cortex volume image [Paxinos et. al. 2012, http://marmosetbrain.org] and aids perspective in the 3D view.

**Figure 7.**
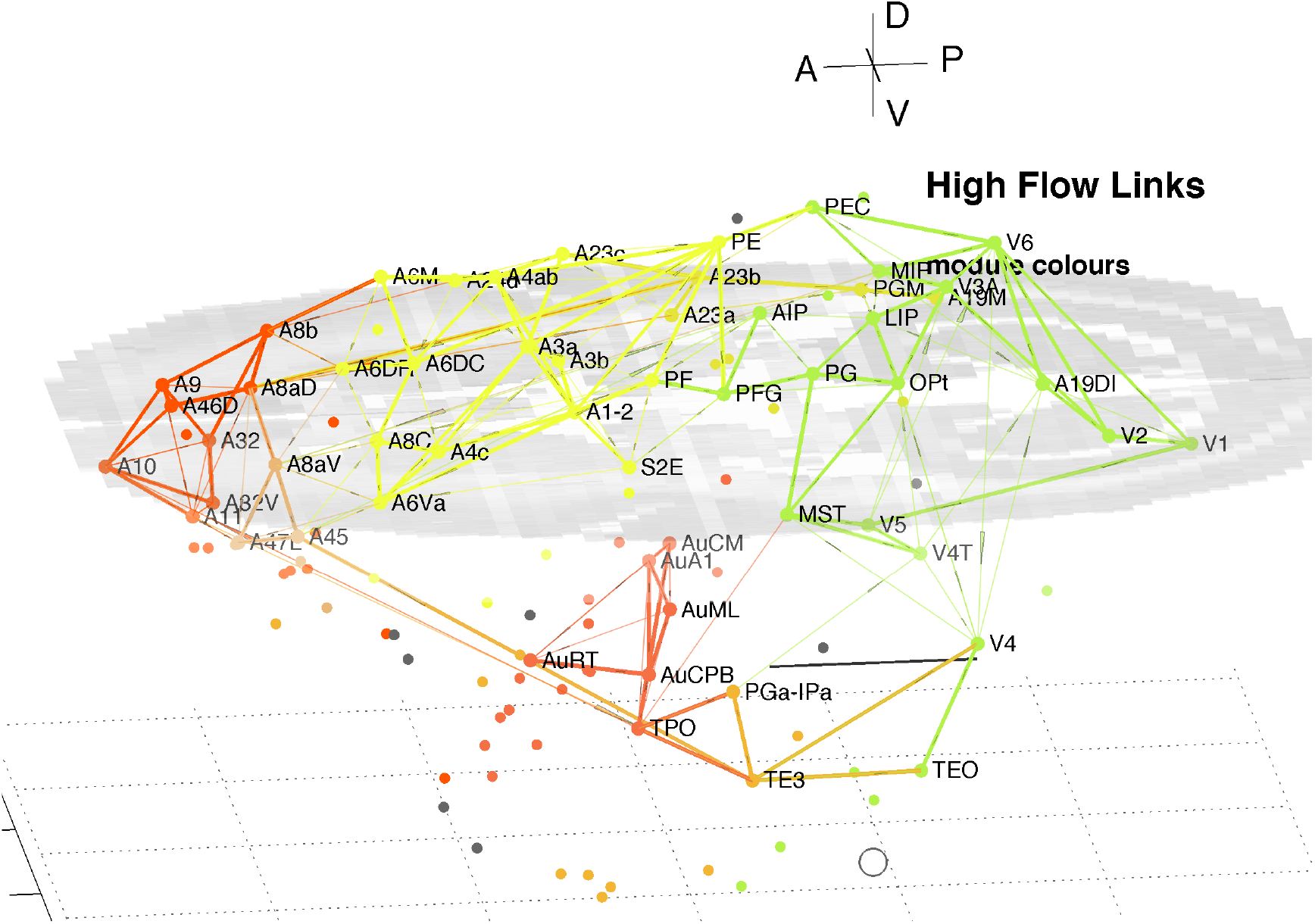
3D plot of all nodes and the top 15% highest probability flow links in the marmoset cortex. Higher traffic links are in bold. Colours identify module membership (cf. Table 1) and follow the marmoset Atlas. The mid level horizontal plane image is a slice from the marmoset cortex volume image to aid perspective, along with the background grid. The A-P scale bar is 5mm.

The clustering evident in Fig. 7 reiterates that found above around hubs and connector nodes and highlights a densely linked cluster in pFC. It is reassuring that methods based on node participation and on link traffic yield similar results. While not conclusive Fig. 7 suggests parallel pathways from sensory areas: from auditory cortex direct to multiple areas of pFC; and from visual cortex via parietal areas to motor areas and pFC. There is some cross coupling of V4 via temporal areas, and of V1, V5 and V6 via MST to auditory cortex. The visual pathways are consistent with a detailed review of the marmoset visual system in [Solomon and Rosa 2014], which noted MST as a transit hub from V6.

## Conclusion

A rescaling of the original marmoset structural connectivity data reveals the underlying in link weights and facilitates a more complete network analysis of the cortical network. The in and out link weights then follow similar distributions (Fig. 1), with rescaled link weights in the range 0.03 – 10^4^. The weight measure is the number of labelled neurons in a target area due to links from a source area, in keeping with the mouse brain data [Oh et. al. 2014, Pailthorpe 2019], but in contrast to the EM measurements of synaptic contacts or areas in mouse retina [Helmstaedter et. al. 2014, Pailthorpe 2016] and fly [Chklovskii et. al. 2010]. While there are a number of options, the relationships between LN and injection and target volumes (Figs. S5, S6) suggest that LNe is a viable measure of link weight. The scaling factor to convert labelled neuron counts to synaptic weights is not clearly defined and warrants further study, thus the interpretation of weight <1 is yet to be clarified.

The distributions of in- and out-weighted degree, ie. node strength, in Fig. 1 suggest that some in links of low-medium weight are missing. A similar plot for mouse brain [Pailthorpe 2019] shows both distributions being quite similar - again suggesting some missing in-weight data here. This is consistent with sampling only 55 source nodes, being the 55, of the 116, anatomical areas used as tracer injection sites. In due course such data should be available and complete the picture emerging here. The link weight – distance relationship, based on the scattered data, marginally follows both an exponential decay, or a power law decay (Fig. S7) consistent with other species.

A decomposition of the network using InfoMap produces 8 modules (Table 1) aligned with dominant cortical regions, and enables the subsequent analysis. Classification of within and between module linkage patterns, using z-scores and participation coefficients, identifies key hubs and connector nodes (Fig. 2). This can be applied to both in and out links. Hubs, with many links between modules or many within a module, provide an anchor for local, densely interlinked clusters (Figs. 3, 4) in pFC, association, auditory and visual areas. By contrast, connector nodes have many links to other modules and are waypoints between the clusters (Figs. 5, 6). Those linkage patterns are explored by following evolving link formation at increasing distances, which reveals the gradual shift from local to global linkage patterns. The analysis reveals PEC, AuRT, A32V and TEO as key in-hubs, with A45, A8C, A1-2, A23c, PF and A11 as possible candidates. A10 is the dominant out-hub, with TE3, AuCPB and AuA1 as candidates also worthy of further investigation.

Another analysis, of cat and macaque hub nodes [Sporns et. al. 2007], used centrality measures and the participation coefficient only, but not the z-score. It was applied to unweighted links and to motifs, which generally are small clusters, typically of 3 nodes. That different methodology identified V4 as a key hub, somewhat consistent with the present study, and A46 (DlpFC) as another key hub. The inclusion of link weights and the z-P classification, used herein, produced hubs in pFC and in sensory, association and motor areas.

A separate, extensive network analysis of the fully connected marmoset subnetwork of 55 nodes ( 55x55 linkage data) [Theodoni et. al. 2022] using the fractional weight measure FLNe produced a hierarchical ordering of areas, from frontal to visual areas. At the top was A6 (motor); AuCPB (auditory); A32, A8aD (frontal cortex); A4c, A6M (motor); and later A11, A46aD, A10 (frontal cortex); continuing through association and somatosensory areas, and on to visual areas at the lowest level. That is somewhat consistent with the ordering in the top 4 modules identified in the present study.

Functional connectivity in marmoset brain, studied by resting state fMRI [Belcher et. al. 2016]. measured as local Functional Connectivity Density, identified 11 hubs across the whole brain. In the cortex: area A24a (Cingulate) was the most prominent, followed by V6, A19M (vis.), A23a,b (posterior Cingulate), and at lowest strength: V1 and V2.

Separately high traffic links across the cortex (Fig. 7) confirm the overall picture presented herein and illustrates parallel auditory and visual pathways. Some features of those paths are consistent with a detailed review of the visual system in marmoset [Solomon and Rosa 2014]. The most significant cluster in pFC comprises 6-8 nodes, two of which (A10, A32V) are dominant hubs with a third (A11) is nearly so. This suggests a central role in the cortical network and warrants further investigation.

The key limitation of this study is that it used incomplete linkage data, with only 55 of 116 areas sampled by tracer injection. One can reasonably expect that those sites were chosen as the most biologically important, so the present analysis produces a partial picture of significant network structures. It extends prior analyses of the marmoset data by including some 1611 links that are missed by trimming the data to 55x55 sources and targets. That excluded 26% of the total link weight measured which now is included in the present study. With the eventual availability of more data that emerging picture can be completed. A separate tracer study of marmoset cortical links [Watakabe et. al. 2023] provides information at columnar level, in a parcellation free analysis, and needs to be integrated with these area level studies. Its analysis will enhance the multiple studies discussed herein.

## Acknowledgements

The author acknowledges valuable suggestions from BD Fulcher and PA Robinson.

## Notes

https://github.com/BrainDynamicsUSYD/MarmosetCortex/

### Competing Interest Statement

The authors have declared no competing interest.

### Summary of Updates

Revision includes more literature review. Results are unchanged.

