## Supplementary material for "Network analysis of marmoset cortical connections reveals pFC and sensory clusters": SupplmMaterial

### Supplementary Data

1. Merged spreadsheet of raw data, attached as file: MarmosetRescale.xlsx .
2. Adjacency matrix derived from LNe, attached as file: MarmosetAdj.csv .
3. Table of data and calculations for the 116 nodes in marmoset cortex, attached as file: Marmoset116node.xls .
Matlab codes at https://github.com/BrainDynamicsUSYD/MarmosetCortex/ .

### Measured Distribution of links and weights, FLNe

The original connectivity data for the marmoset cortex was downloaded [Majka et. al. 2016, http://marmosetbrain.org] as a link list text file. The FLNe values were derived from retrograde viral tracer injections to 55 target areas, and revealed 3474 links from a possible 116 source areas; that is 26% of the possible links in the subsampled full network. This is significantly sparser than the 62.4% fraction of links in the full connected 55x55 subnetwork [Theodoni et. al. 2022]. Use of more injection sites likely will yield a larger fraction of links amongst the 166 nodes, intermediate between those two values. The measured link in-weights were normalised to form the Fraction of Labelled Neurons extrinsic, FLNe, a measure that was introduced for earlier macaque data [Markov et. al. 2011, 2014]. Thus for each node all in-weights sum to 1, so that differences between nodes are factored out, and the weighted in degree of each connected node is 1, or 0 if no links were measured. However this data does preserve the number of links. This is reflected in the plot of in-degree, k-in, as a distribution of the number of links, presented in Figure S1, along with k-out for the out links.

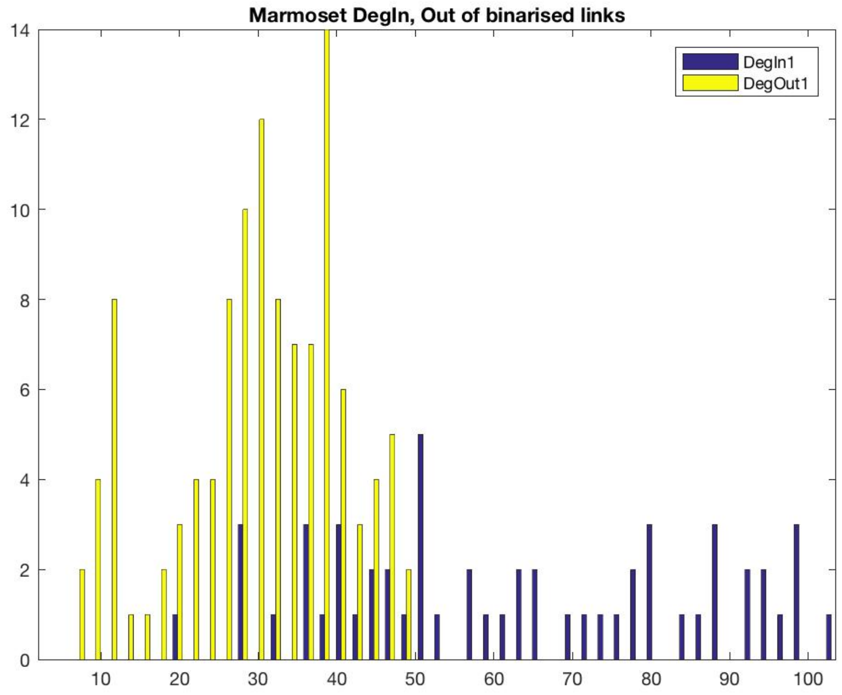

Figure S1. Distribution of the number of in- and out- links (un-weighted degree, k) measured for areas of the marmoset cortex.

The maximum number of out links is 48, much less than the maximum, 104, of in links. The clear asymmetry arises from only 55 target nodes (i.e. tracer injection sites) being measured thus far, from amongst 116 possible sources. There are likely to be some other targets not yet captured in the present dataset. The number of out links is better fit by a normal, rather than log normal, distribution (log likelihood comparison), as discussed below. For the in links the distribution is better fit by an exponential distribution – that may change as more target sites are studied and more data becomes available (note the smaller number of counts). The data set still shows the variation in the number of in links to each node (Fig. S1). Note that the analysis of the fully connected, 55 node subnetwork produces more symmetry between in and out degree: cf. Fig 2b of [Theodoni et. al. 2022].

The weighted degree, or node strength, distributions are shown in Figure S2. The out-degree appears to have an exponential decay, as observed for other species, eg. worm C. Elegans [Varshnay et. al. 2011], mouse retina [Pailthorpe 2014] and mouse brain [Oh et. al. 2014, Pailthorpe 2019]. However for each node the sum of all in-weights is either 0 or 1, a striking result. This follows from FLNe being a fractional measure which sums to either 1 or 0 for each node. Note that a similar analysis of the 55x55 connectivity matrix would produce misleading results since it is formed by deleting the 61 empty columns belonging to the non injection sites. It also requires deletion of the corresponding 61 rows that do contain 1611 non-zero links: these are the out links from the non sampled areas to any of the 55 injection sites. That step excluded 26% of the total link weight measured in the cortex, which now is included in the present study. Those out links (A-B) also happen to be in links (B-A) that were counted in forming FLNe. Those are sampled in Fig S2.

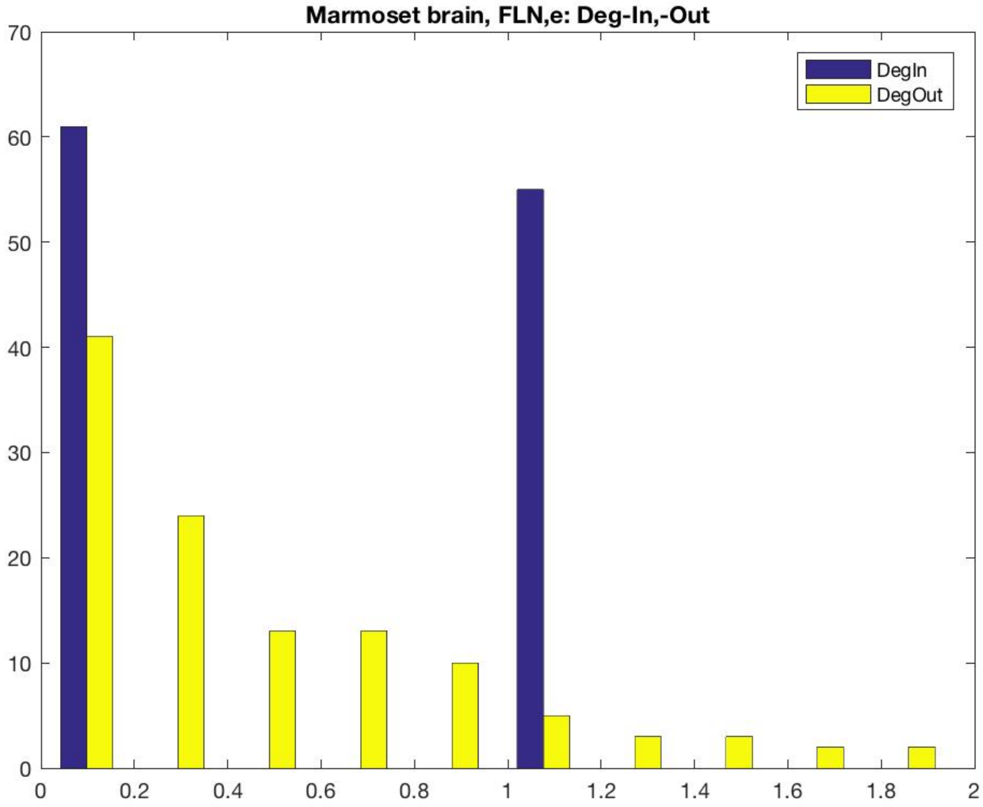


Figure S2. Distribution of the weighted in- and out- degree measured as FLNe for areas of the marmoset cortex.

The out link weight distribution is best fit by an exponential distribution (not shown). The data are more clearly displayed on a $\log_{10}$ scale for clarity as in Figure S3 (the original exponential is so steep that details cannot be judged from the plot). The original normalisation of those fractional weights means that they span the range $4.6 \times 10^{-6}$ - 0.61.

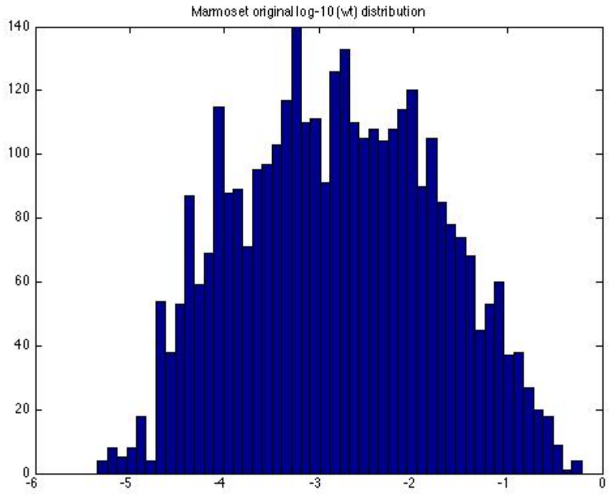


Figure S3. Distribution of all link weights measured by FLNe (counts vs. $\log_{10}$(FLNe)) for anatomical areas of the marmoset cortex.

**Rescaled link weight data, LNe**

The more extensive measurements (Majka et. al. 2020] includes sufficient detail to enable the fractional normalisation, used above, to be recovered, and to produce the underlying link weight. That normalisation factor varies for each target node since it is a measure of the weighted in degree, or node strength. The additional data includes: repeated injection volumes, number of labelled neurons (LN) both intrinsic (to the injected volume) and extrinsic (all source neurons): LNi, LNe, respectively. Note that there is significant variation across the repeat experiments (ie. multiple injections to a target area).

The question of normalisation has also been explored in detail for mouse [Rubinov et. al. 2015] with four measures (CS, NCS, CD, NCD) variously used. They are: Connection Strength (CS): the number of connections from the whole of the source region to the whole of the target Region; Normalized Connection Strength (NCS): the number of connections from one unit volume of the source region to the whole of the target region; Connection Density (CD): the number of connections from the whole of the source region to one unit volume of the target region; and Normalized Connection Density (NCD): the number of connections from one unit volume of the source region to one unit volume of the target region. Such may be contemplated here, to take into account the variations in injected volumes of tracers, or volumes of target or source areas. One might also use an intensive measure such as LNe /$mm^3$ (eg. LNe1), eg. LNe per unit volume injected into the target area, etc. Network analysis of the Marmoset data was also conducted using such measures (not reported). Guidance on which to use is best provided by dependencies revealed in the raw data, or by the research questions being addressed.

Plots of repeat measurements of labelled cells counts, LNe and LNi, vs. injection volume for a single area suggests that LNe/InjVol might be an appropriate measure of connection weight into each target area. However that suggestion is not sustained by detailed statistical analysis, presented below. Now using LNe the resulting link weight distribution ($log_{10}$ scale) is shown in Figure 1, with with the node degree distributions in Fig. 1. Both in- and out-degrees now follow similar distributions, as usually observed in other systems

Next is discussed the details of recovering the normalisation implicit in the fractional weight measure, FLNe. The Supplementary Information of [Malka et. al. 2020], and attached files, contains significant detail of the raw tracer injection data. The six fluorescent tracers described in their Methods section are: FR, FE, FB, DY, and CTB (2 variants). DY labels the cell nucleus while the other 5 label the cytoplasm. They are retrograde tracers, so label the links from source area cells. To proceed two tables needed to be combined manually (in Excel): the first is associated with their Figures 5A, S1 and S2 and also Suppl. Figs. 4c, 4e, 4g. The underlying data is listed in many tabs of a linked Excel data file: 41467_2020_14858_MOESM5_ESM.xlsx. It covers 147 experiments, and for each lists: Case ID, Tracer, Anatomical area, its volume (Vol, $mm^3$) and cell count. The second table is based on their Supplementary Figures 1C, 1D and 1E. For 143 experiments it lists: Case ID, Tracer, Total number of Labelled Neurons Normalized, Number of intrinsic Labelled Neurons Normalized, and Injection Volume (InjVol, $mm^3$). The common 143 experiments were analysed, as described below. The injection coordinates are available separately online at: www.marmosetbrain.org/injection. The injection sites are within previously reported 55 targets.

These two tables can be merged via a pair of common entries: Case ID and Tracer (nb. there is a need to omit 4 experiments: 143 entries have common data). The merged spreadsheet, including detailed working, is attached at Supplementary Data. Two examples illustrate the scope of the data: For target area A10, of volume 25.39 $mm^3$, six injections were made (2 of CT, 2 of DY, FB and FR), ranging in volume from 0.03 to 1.13 $mm^3$, or 0.1 – 5% of the anatomical area. The total number of labelled neurons ranged from 4.7k – 35K. For area V2, of volume 118.89 $mm^3$, nine injections were made (4 x FR, 3 x DY, FB, FE), of volumes 0.02 – 0.56 $mm^3$, or 0.1 – 0.5% of the area. The total number of labelled neurons ranged from 3.8k – 38k. This illustrates significant variability in the extent of sampling, even though a similar and significant number of neurons were labelled in each case. The Supplementary Information in the original report [Majka, et. al. 2020] examined such variability in detail. In the present analysis of repeat experiments a sampling metric, (Vol/ k-In) / InjVol, was also used to judge how well the injected tracer might be able to cover the average volume per link detected. Low values (~ 0.1, 1) indicated strong sampling, while high values (~10, 100) possibly indicated that too little tracer likely was injected to cover all possible links. Generally that was associated with small injection volume and a large number of links detected. The latter case mostly applied to the larger anatomical areas such as V1, V2. Very high values (>50) suggested that a few measurements might be significant outliers that could be omitted from the calculation of mean values. That only applied to one case each in V1 and V2, as annotated in the attached Excel spreadsheet (Supplementary Data). The experimental protocol (inject, wait 3-22 days (median 15)) was designed to allow sufficient time for the tracer to diffuse, and/or be transported, from the small injection volume across the target neurons, then along connected dendrites and axons to the source neurons. Then “signal” was detected by fluorescent imaging of all those cells and comprised counts of labelled neurons, LN, intrinsic to and extrinsic from the injection area. Test calculations confirm that it is reasonable to assume that sufficient tracer diffusion occurred to sample the target area.

**Calculation of link weights from LNe**

The procedure to remove the “fractional” from the originally reported weighted connectivity (or adjacency) matrix, ie. recover LNe from FLNe for source-target pairs, is as follows. The two data tables described above were merged in Excel via the joint common entries of case ID (CJ100 – CJ94) and Tracer. They were then resorted by area acronym (A1-2 to V6), so that repeated measurements of a single area were grouped together. Several options were explored by examining dependencies of LNe and LNi on tracer Injection Volume and Target volume. Thus LNi/InjVol and LNe/InjVol was calculated for each experiment, along with their means for each area. Mean LNi and LNe across repeat experiments were also calculated. Note also that LNtot = LNi + LNe, was recorded in the original data. These data provided the basis for the analysis described below (at Tests of Variability of LNi and LNe) and in the main text. Details of calculations are attached as an Excel file (Supplementary Data).

For source (s) – target (t) pairs one can calculate the underlying link weight LNe(s, t) from the previously reported FLNe(s, t) by using the LNe(t) that are now available with the recent data [Malka et. al. 2020]. The normalisation implicit in the fractional measure FLNe can be seen explicitly as FLNe(s, t) = LNe(s, t)/ LNe(t). That can be reversed by: LNe(s, t) = FLNe(s, t) x LNe(t), which then provides the adjacency (or connectivity) matrix used in the present analysis. This is consistent with eq. 1 of [Majka et. al. 2020] since the LNe already excludes the “neurons identified in area A” (i.e. the target). The data report LNe and LNi separately and for each repeat experiment LNtot = LNe + LNi holds, indicating that intrinsic neurons are already excluded in each case. The mean values of LNtot and LNi were formed (excluding 2 significant outliers, as noted above and annotated in the attached spreadsheet) and used to calculate the mean LNe used to estimate of link weights. This approach is also consistent with the definition proposed originally [Markov et. al. 2011]. The distribution of link weights measured by LNe is shown in Figure S4.

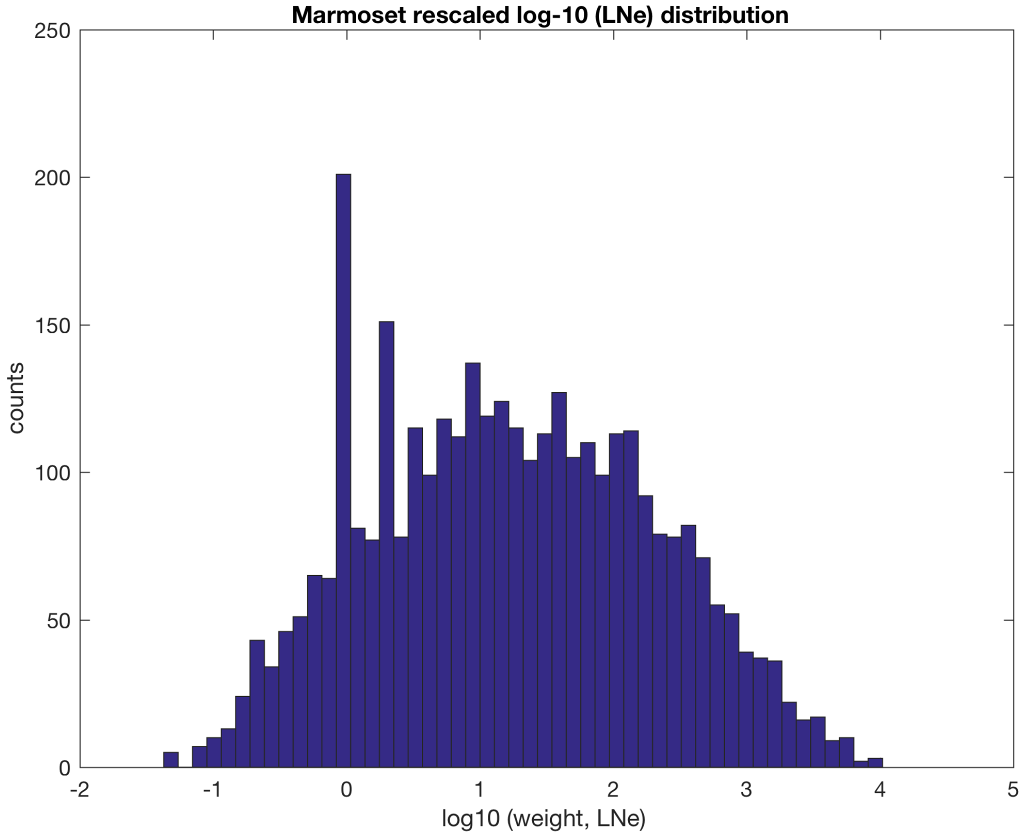


Figure S4. Distribution of rescaled link weights, plotted as $\log_{10}$ LNe, for areas of the marmoset cortex.

In network terms the measured LNe is equal to the weighted In Degree of the injected target node (ie. node strength): this can be calculated both for out links (sum over linked targets) and for in links (sum over linked sources). A more explicit notation, LNe(t) identifies that the injection site for the retrograde tracer is the target node in network terms. This LNe(t) quantifies the totality of incoming links from linked source nodes: i.e. LNe(t) = Σ LNe(s, t), where the sum is over all linked source nodes (s). That sum is just the weighted In Degree of the target node, shown in Fig. 1 (main text). Note that this varies across nodes (areas) while, by contrast, the corresponding fractional measure FLNe(t) is either 0 (no incoming links), or 1 (target linked from one or more sources). The resulting array of link weights, the adjacency matrix, is attached at Supplementary Data. The two evident spikes in Fig S4 are more pronounced than in Fig. S3. They correspond to weights 1 and 2 (ie. $\log_{10}$ (weight) = 0, 0.3), and possibly warrant further investigation. Examination of the connectivity matrix of FLNe values indicate many common entries of those values that might contribute to this effect?

**Tests of Variability of LNi and LNe.**
Plots of labelled cell counts, shown in Figure S5, reveal an approximately linear relationship between detected LNe and injected volume of tracer. Two examples with many repeats (areas V1, V2) are highlighted in Fig. S5 to highlight the approximately linear trend for individual areas. Note that the slope varies for each area, with those points locally scattered. That slope warrants further attention since it may provide clues to the scaling of synaptic weights. The

linear trend suggested that LNe might be an appropriate measure of the extent of connections into each target area.

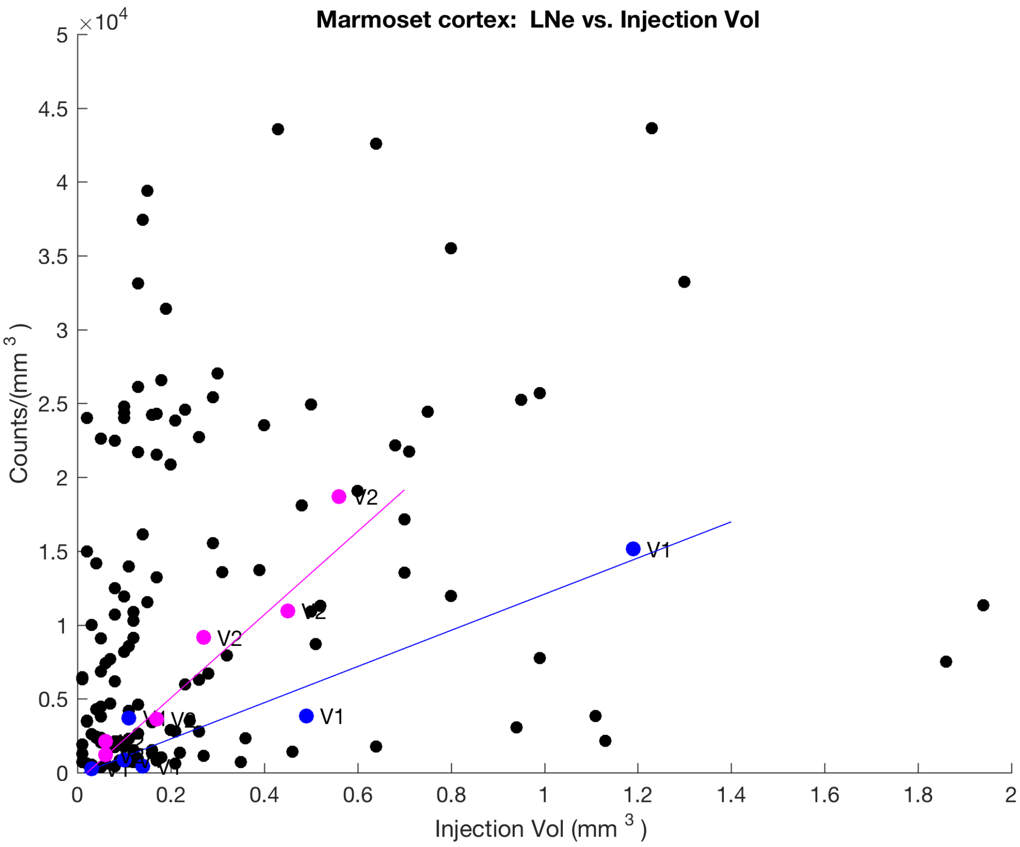


Figure S5. Repeated injections of tracers into target areas: raw counts, LNe vs. volume of tracer injected. Repeats for a single area (eg. V1, blue or V2, magenta) follow an approximately linear trend, with varying slopes as shown by the linear fit lines.

These relationships suggest that cell counts might be normalised by the injection volume to provide a common basis for comparison. Figure S6 shows a plot of intrinsic labelled cell counts to test such a normalisation: LNi/InjVol (injection volume) vs. the volume of the target area (within which the injection sites are located). Repeated measurements appear as a vertical stack in fig. S6, and generally within a tight range. Colours cycle through repeat experiments. Three prominent outliers are: for target areas A46D (experiment case ID CJ800, tracer DY), A4ab (CJ173, tracer DY) and V2 (CJ190, tracer "Dy"). These single measurements, from amongst 2 – 9 repeats, each involved a very small volume (0.01 or 0.02 $mm^3$) of injected tracer, making the measured result more sensitive to the normalisation. For that single V2 case the measured LNi was the largest in the data set; the injection coordinate was the most posterior of those for V2, possibly suggesting the location might be separately delineated? The next two other results are A3a (CJ178, tracer DY) and V4 (CJ182, tracer DY), and appear to be the next most under sampled by tracer. Any designation as possible outliers is not definitive, merely suggestive for further investigation.

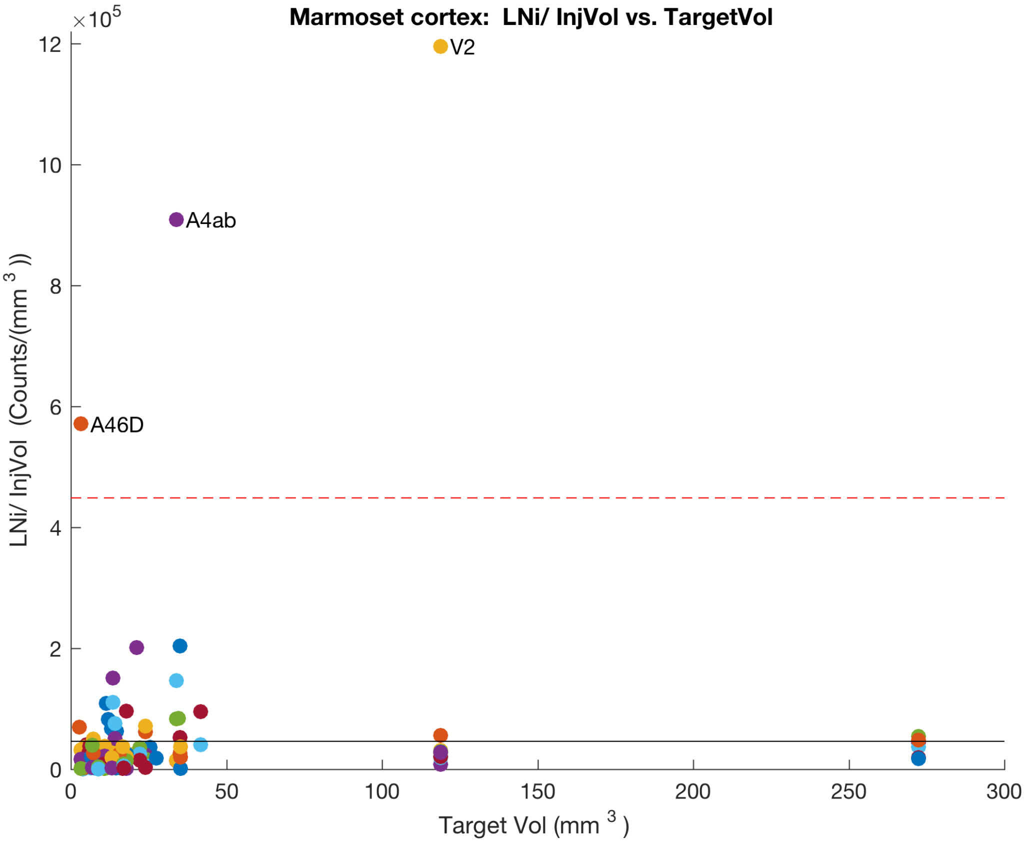


Figure S6. Counts of intrinsic labelled neurons normalised by injected volume of tracer vs. the volume of the target area. Possible outliers (cf. text) are labelled. The mean (solid line) and mean + 3 sd (dashed line) for all nodes are shown. Colours are cycled by Matlab as it scans repeat experiments.

A plot of LNe/InjVol vs. target volume reveals a similar trend to that shown in Fig. S6 –with the same three outliers, plus A3a. Thus a choice of LNe/InjVol as a link weight measure was also considered. However that suggestion is not sustained by detailed statistical analysis, as discussed below. The plots highlight possible outliers amongst the repeated experiments.

These results suggested a possible normalisation for LNe, as mentioned. A log-log plot of LNe vs. InjVol (ie. a re-plot of Fig. S5) suggested a linear trend. However detailed statistical tests revealed that such a linear fit was not significantly better that a model that assumed LNe was constant with Injection Volume: the differences were marginal. A log likelihood (LL) comparison was made, using the Matlab Statistics Toolbox, of two models: (i) LNe = constant (the mean), and (ii) LNe/InjVol = constant, as listed in Table S1, below. Note both cases have the same number of degrees of freedom (142), so the log likelihood comparison is equivalent to the Akaike Information Criterion test. It is clear that model (i) LNe = constant, has the maximum LL.

| Model i): LNe = $1.51 \times 10^4$ | ii) LNe/InjVol = $1.358 \times 10^5$ |
|---|---|
| t statistic: 12.78 | 5.95 |
| p: $2.25 \times 10^{-25}$ | $1.96 \times 10^{-8}$ |
| log likelihood: $-1.569 \times 10^3$ | $-1.992 \times 10^3$ |

Table S1. Statistical comparison of two models: LNe = constant and LNe ~ 1/InjVol.

Despite the variability and trends evident in Fig. S1, statistical analysis does not support use of LNe/InjVol over LNe as a measure of link weight. Thus LNe was adopted herein, to produce the results in the main text. A repeated analysis for a reduced dataset of 55 targets x 55 sources produced similar results. Other tests of internal consistency were examined: eg. plot LNe1 vs k-in (# links): there is an approximately linear trend, with significant scatter.

**Decay of link weights with distance using LNe**

The volume image of the marmoset cortex [Paxinos et. al. 2012, www.marmosetbrain.org] and associated array of voxel labels enable delineation of each anatomical area. The volume of each anatomical area was calculated by counting 3D voxels corresponding to each labelled area in the Marmoset brain atlas [Paxinos 2012]. The calculated centroid, or centre of mass, of those voxels was taken to be the node coordinates, and used to calculate inter-node distances, i.e. the link distances. This is consistent with [Majka et. al. 2020]. Then a plot of link weights, on a $\log_{10}$ scale, vs. distance in Figure S7 indicates an exponential trend. A linear fit (Matlab *cftool*) of $\log_e$(wt) vs. dist yields weight ~ $e^{-dist/4.57(mm)}$ ($R^2$ = 0.16, 95% CI 4.24, -4.93]). A log-log plot also exposes a power law trend and follows weight ~ $dist^{-2.00}$ ($R^2$ = 0.23, 95% CI [-2.12, -1.87]), consistent with other species [Rubinov et. al. 2015]. The large scatter causes the linear fit to be marginal in both cases, slight favouring the power law.

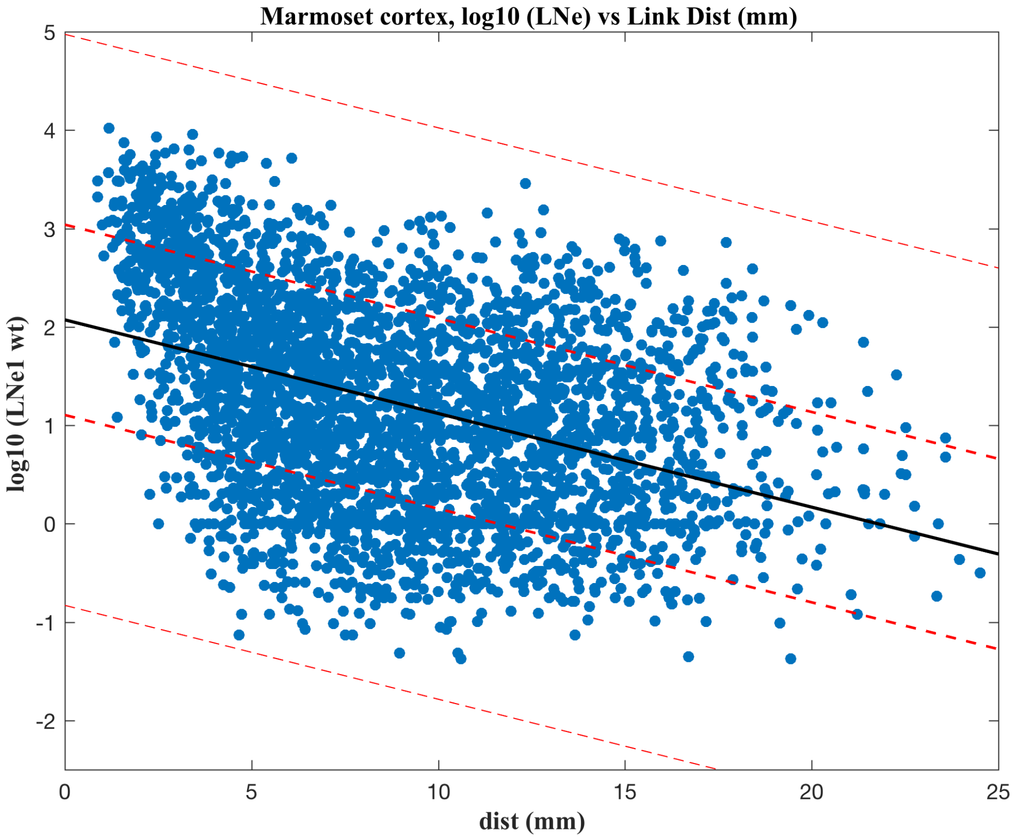


Figure S7. Link weight – distance plot, showing exponential decay of link weights with link length. The $\log_{10}$ scale is more intuitive to read.

Note that all rescaled link weights, while scattered, are within three standard deviations (sd) of the mean fit line (estimated using demeaned data).

**Network modules and hubs derived from FLNe**

The InfoMap modular decomposition calculated using fLNe as a measure of link weights is presented in Table S2.

| Module | Flow in Module | Key areas | Lobes / Regions | Color |
|---|---|---|---|---|
| 1 | 0.326 | A6Va, A4c, A8c, A4ab | Motor, Somato Sensory | yellow |
| 2 | 0.264 | MIP, AIP, Opt, PG | PPC, Vis | green – light |
| 3 | 0.1 | paIM, STR, PaIL, AuRTL | pFC (Dl, M), OrbpFC | Red/ pink |
| 4 | 0.1 | MOB, AON, PIR, ENTl | Aud, Insular Ctx | Salmon |
| 5 | 0.0 | A47L, A47M, A8aV, | pFC (Vl, Dl), OrbFC, Cing/RSP | Green – light |
| 6 | 0.0 | A23a, A23b, A29d, A30 | Cing/RSP | Yellow/ brown |
| 7 | 0.0 | OPAI, AI, DI, TE1 | OrbFC, InsulCtx, LTempCtx, VTempCtx | Green - lime |
| orphans | 0.0 | ProM, Gu, TPPro, APir | VlpFC, OrbFC, Amygd | n/a (gray) |

Table S2. Infomap modules for marmoset cortex ipsi-lateral links, using the FLNe measure of link weights. Key members, in order of probability flow in the module, and color coding follows the Marmoset Atlas and as used in the figures are listed.

Using FLNe the motor - somatosensory system is the top ranked module, somewhat reminiscent of a rodent brain. The orphans have no detected links: possibly arising from their being only 55 target sites for injections, thus many in links may have been missed in the data available so far. The assigned colours follow those used in the Paxinos atlas (available as R G B values) for the dominant nodes in each module. Analysis using LNe is presented in the main text.

The identification of network hubs (cf. Methods) [Guimera et. al. 2005] is depicted by 2D plots of $z_i$ vs. $P_i$, divided into regions that classify nodes and types of hubs, as illustrated in Figure S8. The regions in the 2D plot and the hubs' classifications follow the heuristics developed for metabolic networks [Guimera and Nunes Amaral 2005] to describe classes or roles assigned to network nodes: Role 6, connector hubs (many links between modules); R5, provincial hubs (links preferentially within module); R4, non hub kinless (links across most, or all, modules); R3, non-hub connectors (many links to other modules); R2, non hubs; and R1, ultra peripheral nodes.

For the marmoset cortex data two sets of results are presented herein: first using the original fractional weights FLNe (below; cf. (cf. Figs S2 and S3) and using the rescaled weights LNe (main text). The resulting module memberships are compared in Tables S2 and 1, while the classification plots, of $z_i$ vs. $P_i$, are presented in Figures S8 (cf. Fig 2. for LNe).

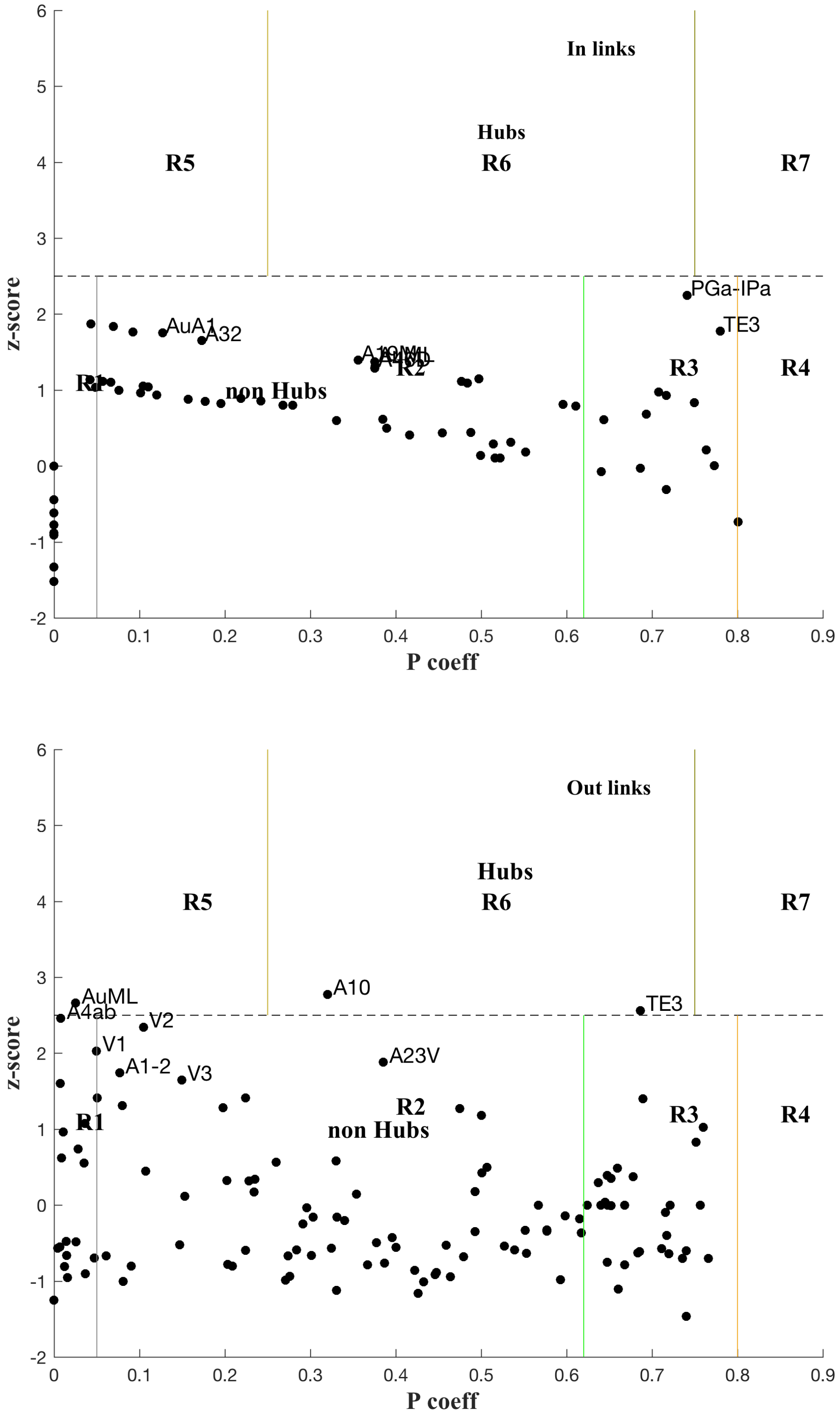


Figure S8. Plot of z-scores and participation coefficients for the marmoset cortex, based on the original link weights, FLNe, for in-links (A. top panel) and out-links (B. bottom panel). Hub nodes and others nearby are labelled.

Using FLNe, where differences were normalised out by using fractional weights, produces no hubs with in-links: this is not surprising since differences between nodes were normalised out in processing the experimental data. Note that PGa-IPa could marginally be classified as R6, rather than R3 as below. For the out-links Connector Hubs (R6) are: A10 (DlpFC) and TE3 (temporal cortex); and Provincial Hubs (R5) are AuML (Auditory cortex) and, marginally, A4ab (SSp). For in-links two non-hub Connector nodes (R3; ie. widely connected across modules) are marginal hubs: PGa-IPa and TE3. Non-hub nodes (R2) that are not far from the hub boundary are AuA1 and A32. Overall these in links do not yield much discrimination between nodes.

For in-links nine nodes are prominent non-hub Connector nodes (R3; ie. widely connected across modules): TE3, A47L, A8aV, A45, TPO, A8b, A8aD, PFG and A6DR - listed in order of decreasing participation coefficient. Three nodes for which in links were measured (ie. an injection site) are classified as peripheral (R1) for in-links: AuCPB, V2 and A4c, along with most (all but 4) nodes that were not measured for in-links, i.e. not injected with tracer. For out links five nodes are prominent non-hub Connector nodes (R3): A47L, A8b, A8aV, PGa-IPa and TE2 - listed in order of decreasing participation coefficient: ie. first listed are most connected to other modules. Another 16 nodes are in the R3 zone, but are less well connected. Seven nodes for which in links were measured are classified as peripheral (R1) for out-links: A32V, along with eight others that were not injection sites: AuAL, AuR, A25, AuRTL, AuRM, TPro and APir. These classifications are listed in detail so that they might be checked to see that they make sense in light of physiological and behavioural observations? Overall there are a significant number of nodes that are well linked across modules or, equivalently, across the cortex.
